# Topology Testing and Demographic Modeling Illuminate a Novel Speciation Pathway in the Greater Caribbean Sea Following the Formation of the Isthmus of Panama

**DOI:** 10.1101/2021.01.22.427733

**Authors:** Benjamin M. Titus, H. Lisle Gibbs, Nuno Simões, Marymegan Daly

## Abstract

Recent genomic analyses have highlighted the prevalence of speciation with gene flow in many taxa and have underscored the importance of accounting for these reticulate evolutionary processes when constructing species trees and generating parameter estimates. This is especially important for deepening our understanding of speciation in the sea where fast moving ocean currents, expanses of deep water, and periodic episodes of sea level rise and fall act as soft and temporary allopatric barriers that facilitate both divergence and secondary contact. Under these conditions, gene flow is not expected to cease completely while contemporary distributions are expected to differ from historical ones. Here we conduct range-wide sampling for Pederson’s cleaner shrimp *(Ancylomenes pedersoni)*, a species complex from the Greater Caribbean that contains three clearly delimited mitochondrial lineages with both allopatric and sympatric distributions. Using mtDNA barcodes and a genomic ddRADseq approach, we combine classic phylogenetic analyses with extensive topology testing and demographic modeling (10 site frequency replicates x 45 evolutionary models x 50 model simulations/replicate = 22,500 simulations) to test species boundaries and reconstruct the evolutionary history of what was expected to be a simple case study. Instead, our results indicate a history of allopatric divergence, secondary contact, introgression, and endemic hybrid speciation driven by the final closure of the Isthmus of Panama and the strengthening of the Gulf Stream Current ~3.5 million years ago. The history of this species complex recovered by model-based methods that allow reticulation differs from that recovered by standard phylogenetic analyses and is unexpected given contemporary distributions. The geologically and biologically meaningful insights gained by our model selection analyses illuminate a novel pathway of species formation that resulted from one of the most biogeographically significant events in Earth’s history.

## Introduction

Tropical coral reefs harbor levels of biodiversity rivaled only by tropical rainforests, yet do so in an environment with few obvious physical barriers to dispersal and gene flow. Although speciation in the sea is not a fundamentally different biological process than on land, the extent to which dispersal and gene flow drive speciation in both ecosystems can work at different scales (Palumbi 1994; Vermeij and Grosberg 2010; Bowen et al. 2013; 2016; Potkamp and Fransen 2019). Most marine species have pelagic larvae that may raft on ocean currents for weeks, connecting distant populations and resulting in large range and effective population sizes (Palumbi 1994; Vermeij and Grosberg 2010; Bowen et al. 2013; 2016; Álvarez-Noriega et al., 2020). Non-allopatric speciation is thus increasingly expected to play a major role in the generation of tropical marine biodiversity (Choat 2006; Bowen et al. 2013; 2016). However, fast moving ocean currents, expanses of deep water, and periodic episodes of sea level rise and fall may act as soft or temporary allopatric barriers that can facilitate both divergence and secondary contact (Quenouille et al. 2011; Cowman et al. 2013). The result is that many reef-dwelling species co-occur with their close relatives in a setting where gene flow may never fully subside, regardless of speciation mode.

That contemporary distributions may mask complex evolutionary histories creates significant hurdles for understanding the origins of tropical marine biodiversity. Complicating matters is that early studies of single loci (primarily mtDNA) could only capture incomplete evolutionary histories, and phylogenetic approaches that employ multi-locus data often concatenate data and/or deal with incongruent gene tree topologies by constructing species trees that model incomplete lineage sorting (reviewed by Degnan and Rosenberg 2009). As evidence continues to build in marine systems for the pervasiveness of speciation with gene flow (e.g. Hurt et al. 2013; Potkamp and Fransen 2019; Simmonds et al. 2019; Titus et al. 2019a; Prada and Hellberg 2020), methods that model reticulate processes are key to testing historical phylogenetic hypotheses and to improving our inferences on the processes that have shaped tropical marine biodiversity.

In the Greater Caribbean (i.e. Caribbean Sea, Bermuda, Florida Reef Tract), Pederson’s cleaner-shrimp *Ancylomene spedersoni* (Chace, 1958; Fig. 1a) is a common member of coral reef communities. An obligate associate of sea anemones, *A. pedersoni* primarily hosts with the corkscrew anemone *Bartholomea annulata* (Briones-Fourzán et al. 2012; Mascaró et al. 2012; Huebner et al. 2019). Together, these species form a hub of ecologically important cleaning stations that are visited by dozens of families of coral reef fishes (e.g. Titus et al. 2015; 2017a; 2019b). Described as a single species throughout its range, molecular studies of mtDNA barcodes have revealed two sympatric lineages along the Florida Reef Tract-one thought to be endemic and one thought to be widespread in the Caribbean- and a third endemic lineage in Bermuda (Titus and Daly 2015, 2017; Titus et al. 2017b). Based solely on a mtDNA phylogenetic analysis with a (Florida, (Caribbean, Bermuda)) topology, Titus et al. (2017b) hypothesized that a chance long distance dispersal event from the Caribbean lineage gave rise to the Bermudan endemic, but were unable to make informed hypotheses regarding the origin of the Florida/Caribbean divergence or whether gene flow has been important during the diversification of the clade.

**Figure 1.**
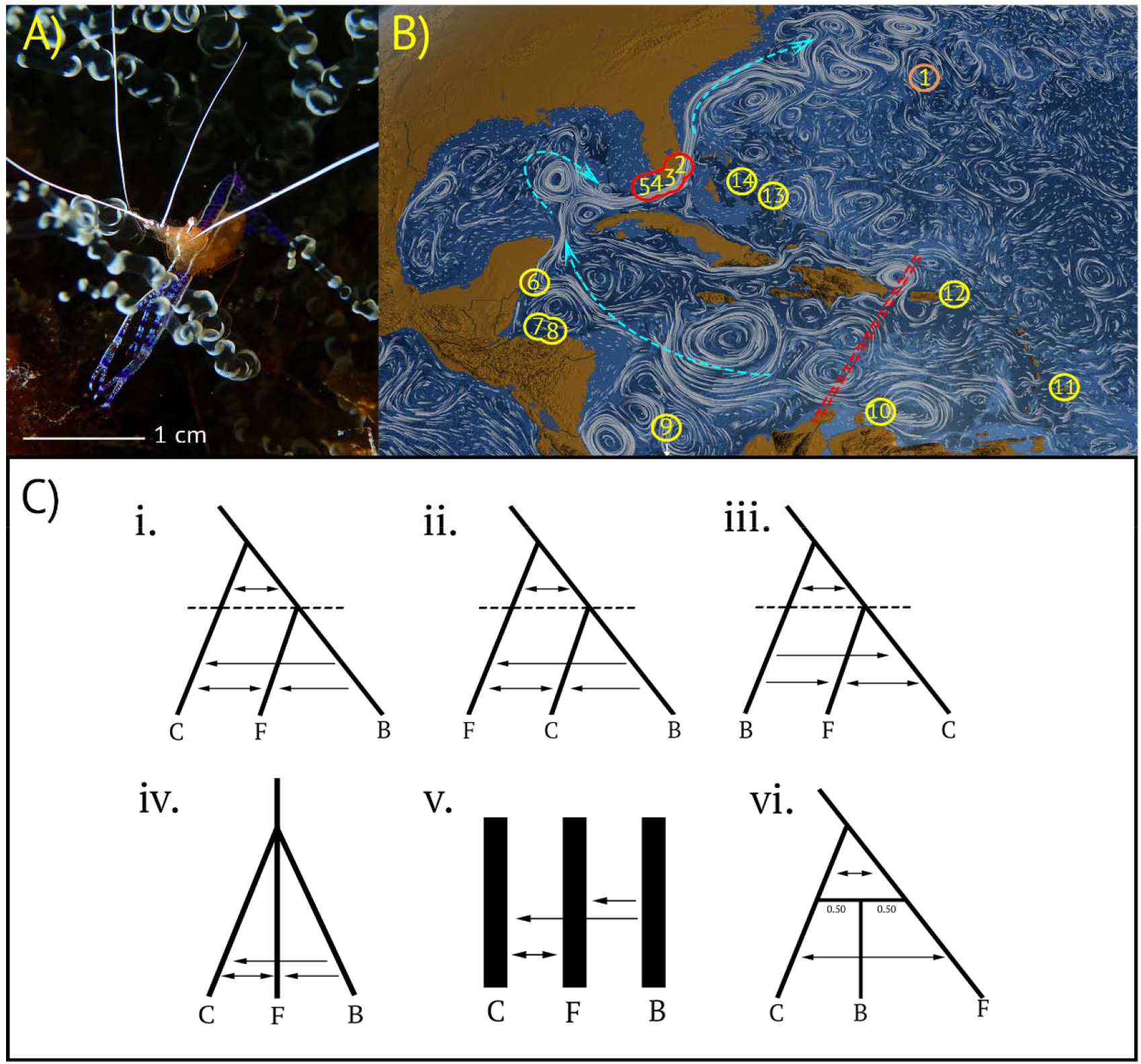
A) Representative image of *Ancylomene spedersoni.* B) Field sampling localities in the Tropical Western Atlantic 1) Bermuda 2) Ft. Lauderdale, FL, USA, 3) Upper Keys, FL, USA, 4) Middle Keys, FL, USA, 5) Lower Keys, FL, USA, 6) Mahahual, Mexico, 7) Utila, Honduras, 8) Cayos Cochinos, Honduras, 9) Bocas del Toro, Panama, 10) Curacao, 11) Barbados, 12) St. Thomas, US Virgin Islands, 13) San Salvador, Bahamas, 14) Eleuthera, Bahamas. Field site circle color indicates geographic range of each putative speces: orange = endemic Bermudan lineage, red = endemic Floridian plus Caribbean lineages, and yellow = Caribbean lineage. Ocean currents depicted by white stylized lines. Current strength corresponds to white line density and length. Directionality of major ocean currents depicted by light blue dashed lines. Map background and ocean currents visualization from NASA/Goddard Space Flight Center Scientific Visualization Studio. Ocean currents visualized from current data collected from June 2005 - December 2007. Double red dashed line demarcates the major phylogeographic break in the Caribbean. C) Examples of the most parameter rich models used in *fastsimcoal2* simulations. i-iii) Isolation-migration models on a fixed species tree topology with variation in the timing and directionality of migration. iv) Isolation-migration models with simultaneous divergence between putative species. v) Island models. vi) Hybrid speciation model where the Bermudan lineage represents a 50/50 hybrid between Caribbean and Floridian species. Hybrid models were built with and without ancestral and contemporary gene migration. C = Caribbean, F = Florida, B = Bermuda. Arrows represent migrations events in the coalescent (i.e. backwards in time). In total, 45 models were built and simulated by considering all pairwise combinations of gene flow between species (see main text)

Here we conduct comprehensive range-wide sampling for *A. pedersoni* and use mtDNA loci and genome-wide approaches to delimit species and disentangle the evolutionary history of this species complex. We combine classic phylogenetic approaches with model-based simulation methods to conduct topology testing and make demographic inferences that allow us to evaluate models that invoke different modes of speciation. We find that regardless of dataset, classic phylogenetic approaches combined with contemporary distributions fail to fully explain the history of *A. pedersoni.* In contrast, our model-based approach highlights the complex role gene flow and hybridization have played in the formation of these species and suggest that historical and contemporary distributions differ. The geologically and biologically meaningful insight provided by our model-based analyses ultimately allow us to identify a novel diversification pathway in the Greater Caribbean following the final formation of the Isthmus of Panama.

## Materials and Methods

### Sampling and DNA Extraction

Whole individuals of *A. pedersoni* were collected from sea anemones by hand, using SCUBA, from 14 coral reef field sites in the Greater Caribbean representing the entire geographic range of the species complex (Fig. 1b; Supplementary Table S1 available on Dryad). A total of 451 individuals were collected and included in this study. DNA was extracted from each individual using QIAGEN DNeasy Blood & Tissue Kits, and total genomic DNA was quantified (ng/μL) using a Qubit 2.0 fluorometer (see Supplementary Methods available on Dryad).

### Sequencing and Dataset Assembly

For each individual we PCR amplified and sequenced a ~650bp long fragment of mtDNA from the cytochrome *c* oxidase subunit I (COI) DNA barcode (see Supplementary Methods). Sequences that were newly generated for this study were deposited into GenBank (Table S1). Following DNA barcoding we sequenced a subset of the 451 individuals using a double digest restriction site associated DNA sequencing approach (ddRADseq). In total our ddRADseq libraries included ~10-15 individuals per field locality (N = 151 individuals; see Supplementary Methods; Table S2 available on Dryad). ddRADseq libraries were prepared using *Eco-*RI-HF and *PstI-HF* enzymes, Illumina compatible barcodes, and a 400-800 base pair size selection range. Two individuals of the closely related cleaner shrimp *Periclimenes yucatanicus* were prepared and included in ddRADseq libraries as outgroup samples. Libraries were pooled and sequenced across eight Illumina HiSeq 2500 lanes using single-end 100bp sequencing.

mtDNA barcode sequence data were assembled using Sequencher 4.9 (Gene Codes) and aligned using MUSCLE (Edgar 2004) in Geneious v10.2.3 (Kearse et al. 2012). Previously published COI sequences from *P. yucatanicus, P. rathbunae, P. crinoidalis, P. perryae*, and *P. patae* were retrieved from GenBank included as outgroup taxa (see Supplementary Methods).

Raw ddRADseq reads were demultiplexed, aligned, and assembled *de novo* using the program pyRAD v3.0.66 (Eaton 2014) as there is no reference genome for *A. pedersoni.* Briefly, we set the clustering threshold (Wclust) to 0.90 to assemble reads into loci and a minimum coverage depth of seven to call a locus (Mindepth). We required an individual sample to contain at least 10,000 consensus loci after clustering, and a locus to be present in 75% of all individuals to be retained in the final dataset. When more than one SNP was present at a locus, only one was randomly selected to create the final unlinked SNP dataset. Because different downstream analyses may or may not require outgroups, and because individual variation in sequencing coverage can influence the number of recovered SNPs, we created three ddRADseq datasets: 1) Full *A. pedersoni* dataset with no outgroup taxa, 2) Reduced *A. pedersoni* dataset with outgroup taxa, and 3) Reduced *A. pedersoni* dataset with no outgroup taxa (see Supplementary Methods and Supplementary Table S2 on Dryad). Raw ddRADseq data were deposited at the NCBI Sequence Read Archive (SRA) under BioProject PRJNA693691.

### Species Delimitation and Phylogenetic Analyses

To test whether Titus et al. (2017b) had identified all *A. pedersoni* species-level diversity throughout the Greater Caribbean, we conducted species discovery and delimitation analyses for both single locus and ddRADseq datasets. For COI DNA barcodes, we used two species discovery approaches: 1) pairwise sequence divergence using the program Automatic Barcode Gap Discovery (ABGD; Puillandre et al. 2012), and 2) statistical parsimony networks using the program TCS v2.1 (Clement et al. 2000; see Supplementary Methods). For the full ddRADseq dataset (Dataset 1), we searched for major genetic partitions in our data with genetic cluster analyses using discriminate analyses of principle components (DAPC; Jombart et al. 2010) in the *adegenet* package (Jombart and Ahmed 2011) in R v3.5.0 (R Core Team 2015). Next, we used Bayes Factor Delimitation* (BFD*; Leache et al. 2014a) to test competing species delimitation hypotheses using species trees estimated in SNAPP (Bryant et al. 2012). Specifically, we tested the current taxonomy of *A. pedersoni* (i.e. one widespread species) versus species delimitations recovered by single and multi-locus species discovery analyses (i.e. two and three species models respectively; see Supplementary Methods).

Phylogenetic analyses were then conducted to reconstruct evolutionary relationships among *A. pedersoni* lineages. Using single locus COI data, we used RAxML 8.9.2 (Stamatkis 2014) and BEAST 1.8.2 (Drummond et al. 2007) to conduct maximum likelihood (ML) and Bayesian phylogenetic analyses (see Supplementary Methods). Next, we used the reduced ddRADseq dataset with outgroup taxa (Dataset 2) to conduct species tree analyses using SNAPP and SVDquartets (Chifman and Kubatko 2014). For both, individuals were assigned to species based on the results of our species delimitation analyses (see Results). Our species tree analysis was run for 1,000,000 MCMC generations in SNAPP, and trees were sampled from the posterior distribution every 100 generations, 10% of which were discarded as burn-in. For SVDquartets, species tree analyses were conducted in PAUP v4.0 (Swofford 2003) using the multispecies coalescent model, all possible quartet combinations, and 1000 bootstrap replicates (see Supplementary Methods).

### Topology testing and demographic model selection

Historical and contemporary gene flow can affect species tree topologies and parameter estimations (e.g., Leache et al. 2014b; Solis-Lemus et al. 2016; Jiao et al. 2020; Rannala et al. 2020), but species tree analyses that use the multi-species coalescent model assume gene tree discordance is the result of incomplete lineage sorting (ILS) rather than gene flow. Our mitochondrial gene trees and ddRADseq species trees produced conflicting, yet fully supported, tree topologies, and genetic clustering analyses suggest ongoing introgression where Floridian and Caribbean *A. pedersoni* lineages co-occur along the Florida Reef Tract (see Results). To account for gene flow and the potential for this to bias our species tree reconstruction and parameter estimates, we incorporated migration parameters into topology testing and demographic model selection analyses to provide statistically supported inference on the evolutionary scenarios that may have given rise to the *A. pedersoni* species complex.

Using the reduced ddRADseq dataset without outgroup taxa (Dataset 3), the multidimensional site frequency spectrum (mSFS), and coalescent simulations in *fastsimcoal2* (FSC2; Excoffier et al. 2013), we built a comprehensive set of 45 evolutionary models across all possible species tree topologies that vary in the timing and directionality of migration between putative species (Fig. 1c; Supplementary Fig. S1 on Dryad). We built twelve models for each possible species tree topology: (Caribbean, (Florida, Bermuda)), (Florida, (Caribbean, Bermuda)), and (Bermuda, (Florida, Caribbean)). Models tested for contemporary and ancestral gene flow between all putative species. We also tested an additional nine evolutionary models that are not explicitly reliant on hierarchical species tree topologies: three island models testing all migration combinations, three models with and without migration where the Caribbean, Florida, and Bermudan species diverged simultaneously forming an unresolved polytomy, and three hybrid speciation models, with and without migration, where both the Caribbean and Florida species provided 50% of the genetic material that gave rise to the endemic Bermudan species (Fig. 1c; Supplementary Fig. S1 on Dryad).

To perform model selection analyses, we constructed a 3-population joint-folded mSFS for putative *A. pedersoni* lineages from Florida, Caribbean, and Bermuda (see Supplementary Methods). To deal with missing data and not violate the assumptions of the mSFS we conducted a down sampling procedure following Smith et al. (2017; see Supplementary Methods). Ten mSFS replicates were built following this approach to account for possible variation in the down sampling procedure and to calculate confidence intervals for parameter estimates. We used a mutation rate of 3.9e^−9^ substitutions per site per generation, calculated from the genome of the crustacean species *Daphnia pulex* (Order Cladocera; Keith et al. 2016), to convert parameter estimate values to years, effective population sizes, and per generation migration rates. Parameter estimates were then further scaled by assuming 2 generations per year because *A. pedersoni* reaches sexual maturity at ~6 months (Gilipin & Chadwick 2017). All model simulations were repeated 50 times for each mSFS replicate (45 models x 10 mSFS replicates x 50 simulations/model/mSFS replicate = 22,500 simulations). From each of the 50 runs, the run with the highest composite likelihood was selected for parameter estimation and model selection. The model with the best fit to the data was selected using Akaike Information Criterion (AIC) and model probabilities calculated following Burnham and Anderson (2002). Parameter estimates and 95% confidence intervals were then calculated for the best fit model. All analyses were conducted on the Owens cluster at the Ohio Supercomputer Center (http://osc.edu).

Finally, we conducted analysis to independently confirm model selection inferences about introgression and hybridization. Specifically, we used ABBA/BABA tests to confirm introgression between Floridian and Caribbean lineages along the Florida Reef Tract, and we conducted hybrid detection analyses using the program HyDe (Blischak et al. 2018) to confirm that the Bermudan endemic lineage received substantial genomic contributions from both Floridian and Caribbean lineages (see Supplementary Methods).

## Results

### Sequencing and Dataset Assembly

After sequencing, alignment, and trimming, we obtained 584 bp of mtDNA *COI* sequence data across 451 individuals (final alignment available on Dryad). ddRADseq library preparation and sequencing resulted in a total of 300.6 million sequence reads across 153 individuals (151 *A. pedersoni* plus two *P. yucatanicus;* Table S2 on Dryad), 276.1 million of which passed quality control filtering and were retained to create the final datasets. Of the 151 *A. pedersoni* sequenced, 16 were not retained in the final dataset because they did not yield at least 10,000 consensus sequences after clustering (Table S2 on Dryad). Requiring a locus to be present in 75% of all individuals resulted in final datasets of varying sizes: 1) Full *A. pedersoni* dataset with no outgroup taxa (N = 135 individuals; 1232 SNPs), 2) Reduced *A. pedersoni* dataset with outgroup taxa (N = 45 individuals; 2101 SNPs), and 3) Reduced *A. pedersoni* dataset with no outgroup taxa (N = 43 individuals; 4673 SNPs, see Supplementary Methods and Supplementary Table S2 on Dryad).

### Species Delimitation and Phylogenetic Analyses

Both single locus species delimitation approaches delimit three *A. pedersoni* lineages (Fig 2a). No new COI lineages were recovered using range-wide sampling, although the Caribbean lineage does show the classic East/West phylogeographic structure at the Mona Passage (Fig 2a). Phylogenetic analyses in RAxML and BEAST recover the same fully supported mtDNA tree topology (Florida, (Caribbean, Bermuda)) as Titus et al. (2017; Fig 2a).

**Figure 2.**
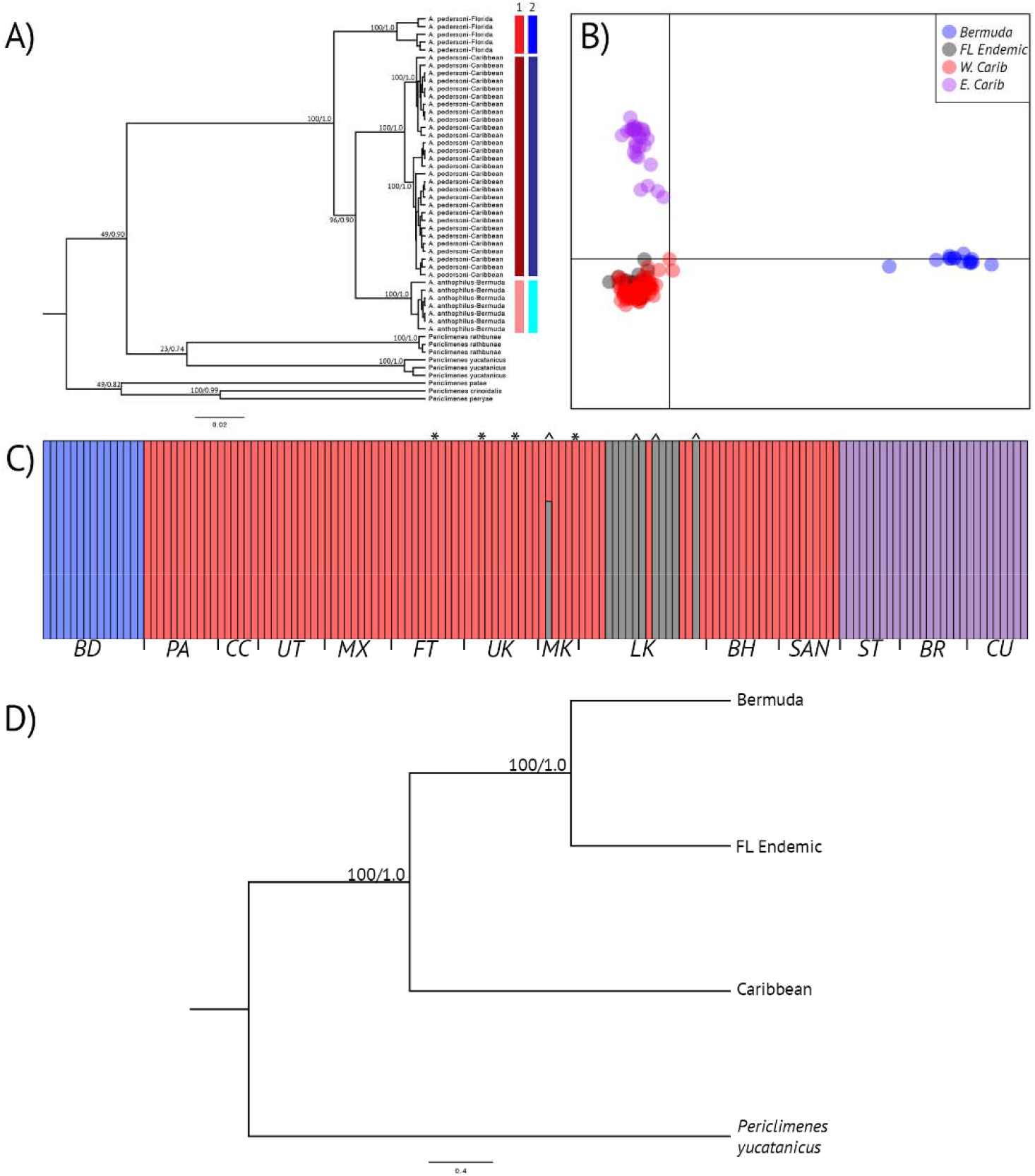
A) Results from cytochrome *c* oxidase subunit *(COI)* phylogenetic gene tree reconstruction *of Ancylomenes pedersoni.* Node labels represent bootstrap and posterior probability values from RAxML and BEAST. Colored bars represent single locus species delimitation results from 1) Automatic barcode gap detection (ABGD) and 2) Parsimony haplotype network reconstruction (TCS). B) Scatterplot depicting K = 4 genetic clusters for *A. pedresoni* throughout the Tropical Western Atlantic using double digest Restriction Site Associated DNA sequencing (ddRADseq). The best clustering scheme was determined by discriminant analysis of principal components (DAPC) and Bayesian Information Criterion. C) Bar plot depicting K = 4 genetic clusters for *A. pedersoni* ddRADseq data using DAPC analyses: Bermuda (blue), Florida Endemic (Gray), Western Caribbean (Red), and Eastern Caribbean (purple). * denotes introgressed individuals with FL endemic COI haplotypes. ^ denotes introgressed inviduals with Western Caribbean COI haplotypes. Sample locality codes are as follows: BD (Bermuda), PA (Bocas del Toro, Panama), CC (Cayos Cochinos, Honduras), UT (Utila, Honduras), MX (Mahauhal, Mexico), FT (Ft. Lauderdale, FL, USA), UK (Upper Keys, FL, USA), MK (Middle Keys, FL, USA), LK (Lower Keys, FL, USA), BH (Eleuthera, Bahamas), SAN (San Salvador, Bahamas), ST (St. Thomas, US Virgin Islands), BR (Barbados), CU (Curacao, Netherlands Antilles). D) ddRADseq species tree reconstruction of the three A. pedersoni lineages as delimited by BFD*. Node values represent bootstrap values andposterior probabilities as determine by SVDquartets and SNAPP.

Using the full ddRADseq dataset, genetic clustering analyses selected *K* = 4 as the optimum partitioning scheme (Figs 2b & 2c). These clusters correspond to the species delimited by COI single locus analyses: Bermuda, Florida, and Caribbean. The DAPC analyses further partitioned the Caribbean species into East and West clusters, reflecting the same intraspecific structuring recovered by the COI analyses. Genetic cluster analyses also demonstrate possible introgression between sympatric COI lineages along the Florida Reef Tract (Fig. 2c). Four individuals with COI haplotypes belonging to the Floridian Endemic lineage cluster with the Western Caribbean *A. pedersoni* lineage, while four individuals belonging to the Caribbean COI haplotype lineage cluster with the Floridian Endemic lineage (Fig. 2c). No individuals outside of Florida differed in their COI and RADseq cluster groupings. Like the COI data, BFD* species delimitation analyses picked the three species model as the best fit to the data (Table S4).

Although the species delimitation analyses fully agree with each other, the phylogenetic analyses between single-locus and genomic data do not. Using species delimitation assignments supported by BFD*, species tree analyses in both SNAPP and SVDquartets recovered fully supported species tree topologies of (Caribbean, (Florida, Bermuda)) that are at odds with the COI phylogenetic tree (Fig 2d).

### Topology testing and demographic model selection

Demographic simulations in FSC2 and AIC model selection placed all model support on a (Caribbean, (Florida, Bermuda)) tree topology, and a demographic history that indicates ancestral divergence occurred in isolation, followed by contemporary introgression between sympatric Caribbean and Florida species, and unidirectional contemporary gene flow from both Caribbean and Florida species to Bermuda (Table 1; Fig. 3). Divergence time estimates place ancestral divergence between the Caribbean lineage and the MRCA of Florida and Bermuda at 3.52 mya (95% CI = 3.02-4.03 mya; Supplementary Table S5), and divergence between Florida and Bermuda at 0.20 mya (95% CI = 0.18-0.22 mya; Table S5). Effective population size estimates were an order of magnitude larger in the Floridian species than for the Caribbean species (Supplementary Table S5). Effective population sizes estimates were less than 1,000 individuals for the putative Bermuda species (Supplementary Table S5). Per generation migration rate estimates from both putative Caribbean and Florida species to Bermuda were low (Supplementary Table S5).

**Table 1.** Akaike Information Criterion results for models simulated in FSC2 for the *Ancylomenes pedersoni* species complex. Model name refers to variants of those depicted and described in Figure 2. *k* = number of parameters in the model, AIC = Akaike Information Criterion, Δ_I_ = change in AIC scores, and *W_i_* = Akaike weights. Prefixes on model names refer to variation in model type and tree topology: A = ((Caribbean, (Florida, Bermuda)); B = ((Florida, (Caribbean, Bermuda)); C = ((Bermuda, (Caribbean, Florida)); Isl = Island Model; Hyb = Hybridization model; Sim = simultaneous divergence model. Models are listed according to their AIC rank and the highest ranked model is highlighted.

| Model | $k$ | Log (P) | AIC | $\Delta_i$ | Model Likelihood | $w_i$ |
| --- | --- | --- | --- | --- | --- | --- |
| i-Mod10 | 11 | -22603.84 | 45229.67 | 0 | 1 | 1 |
| ii-Mod6 | 10 | -22676.34 | 45372.69 | 143.01 | 8.8e-32 | 8.8e-32 |
| ii-Mod10 | 11 | -22752.57 | 45527.15 | 297.48 | 2.5e-65 | 2.5e-65 |
| iii-Mod11 | 14 | -22999.99 | 46027.99 | 798.32 | 4.4e-174 | 4.4e-174 |
| i-Mod11 | 12 | -23010.73 | 46045.46 | 815.79 | 7.1e-178 | 7.1e-178 |
| ii-Mod5 | 11 | -23208.93 | 46439.85 | 1210.17 | 1.6e-263 | 1.6e-263 |
| ii-Mod9 | 12 | -23219.90 | 46463.79 | 1234.11 | 1.0e-268 | 1.0e-268 |
| ii-Mod11 | 12 | -23288.88 | 46601.75 | 1372.08 | 1.1e-298 | 1.1e-298 |
| Sim-Mod3 | 9 | -23311.22 | 46640.44 | 1410.76 | 4.5e-307 | 4.5e-307 |
| ii-Mod7 | 9 | -23322.72 | 46663.43 | 1433.75 | 0 | 0 |
| iii-Mod7 | 11 | -23331.81 | 46685.61 | 1455.93 | 0 | 0 |
| iii-Mod6 | 11 | -23350.94 | 46723.87 | 1494.19 | 0 | 0 |
| ii-Mod3 | 10 | -23421.51 | 46863.02 | 1633.34 | 0 | 0 |
| ii-Mod2 | 9 | -23503.47 | 47024.93 | 1795.25 | 0 | 0 |
| ii-Mod4 | 10 | -23510.66 | 47041.32 | 1811.64 | 0 | 0 |
| Isl-Mod3 | 6 | -23567.15 | 47146.29 | 1916.61 | 0 | 0 |
| iii-Mod10 | 8 | -23617.00 | 47249.99 | 2020.31 | 0 | 0 |
| Isl-Mod2 | 7 | -23625.89 | 47265.79 | 2036.11 | 0 | 0 |
| iii-Mod9 | 9 | -23681.36 | 47380.72 | 2151.04 | 0 | 0 |
| ii-Mod8 | 8 | -23742.41 | 47500.83 | 2271.14 | 0 | 0 |
| i-Mod12 | 9 | -23742.79 | 47543.59 | 2313.90 | 0 | 0 |
| ii-Mod12 | 9 | -23796.70 | 47611.40 | 2381.72 | 0 | 0 |
| Hyb-Mod2 | 10 | -23804.42 | 47628.84 | 2399.16 | 0 | 0 |
| iii-Mod8 | 9 | -23821.71 | 47661.43 | 2431.75 | 0 | 0 |
| ii-Mod1 | 7 | -23910.76 | 47835.53 | 2605.85 | 0 | 0 |
| Hyb-Mod3 | 11 | -24004.63 | 48031.26 | 2801.58 | 0 | 0 |
| Sim-Mod2 | 7 | -24593.07 | 49200.14 | 3970.46 | 0 | 0 |
| Hyb-Mod1 | 8 | -24598.03 | 49212.06 | 3982.38 | 0 | 0 |
| iii-Mod3 | 10 | -24691.93 | 49403.87 | 4174.19 | 0 | 0 |
| Sim-Mod1 | 5 | -24723.30 | 49456.61 | 4226.93 | 0 | 0 |
| iii-Mod5 | 10 | -24742.51 | 49505.02 | 4275.34 | 0 | 0 |
| i-Mod7 | 9 | -24766.59 | 49551.18 | 4321.50 | 0 | 0 |
| i-Mod4 | 10 | -24773.07 | 49566.14 | 4336.46 | 0 | 0 |
| i-Mod5 | 11 | -24789.54 | 49601.07 | 4371.39 | 0 | 0 |
| i-Mod9 | 12 | -24792.60 | 49609.20 | 4379.52 | 0 | 0 |
| i-Mod2 | 9 | -24796.43 | 49610.87 | 4381.19 | 0 | 0 |
| iii-Mod4 | 9 | -24807.98 | 49633.96 | 4404.28 | 0 | 0 |
| iii-Mod2 | 8 | -24809.91 | 49635.82 | 4406.14 | 0 | 0 |
| i-Mod3 | 10 | -24809.13 | 49638.27 | 4408.59 | 0 | 0 |
| i-Mod8 | 8 | -24816.92 | 49638.27 | 4420.16 | 0 | 0 |
| iii-Mod12 | 8 | -24825.62 | 49667.25 | 4437.57 | 0 | 0 |
| i-Mod6 | 10 | -24832.44 | 49684.87 | 4455.19 | 0 | 0 |
| iii-Mod1 | 7 | -24856.22 | 49726.44 | 4496.76 | 0 | 0 |
| i-Mod1 | 7 | -24919.15 | 49852.29 | 4622.61 | 0 | 0 |
| Isl-Mod1 | 6 | -25386.13 | 50784.27 | 5554.59 | 0 | 0 |

ABBA/BABA tests confirm that shared alleles between Floridian and Caribbean species found along the Florida Reef Tract are the result of introgression rather than incomplete lineage sorting (Supplementary Table S6). Hybrid detection analyses in HyDe indicate that the Bermudan species has a significant signature of hybridization between putative Florida and Caribbean species at both the population and individual level (Supplementary Table S7). HyDe estimates that ~48% of the Bermudan species genome comes from the Caribbean *A. pedersoni* lineage (Supplementary Table S7).

## Discussion

Our evolutionary reconstruction of *Ancylomenes pedersoni* highlights the complexity of the processes that can generate species-level biodiversity. While our topology tests ultimately agreed with our ddRADseq species tree topology, the demographic insights gained from this approach underscores the importance of using model-based analyses that account for gene flow to make improved inferences beyond what is possible with classic phylogenetic methods alone. Based on our best fit demographic model (Fig. 4), the Floridian and Caribbean *A. pedersoni* lineages diverged ~3.5 mya without gene flow. This divergence time coincides with the final closure of the Isthmus of Panama (IOP) and strengthening of the Gulf Stream, a high-volume and high-velocity ocean current that originates in the Florida Straits and separates the Florida Reef Tract from the Caribbean Sea (Fig. 1; Huag and Tiedemann 1998; O’Dea et al. 2016). Combined with a demographic signature of introgression along the Florida Reef Tract occurring more recently in the evolutionary past, we interpret the initial divergence between these lineages to have occurred allopatrically across the Florida Straits, with secondary contact coming later when the Caribbean lineage was able to disperse back across this barrier. Ultimately, the Gulf Stream facilitated divergence and dispersal in *A. pedersoni*, and carried pelagic larvae from both Floridian and Caribbean lineages >1500km to Bermuda where they hybridized and remained isolated enough to warrant status as an endemic hybrid species. The evolutionary history of *A. pedersoni* is thus far more complex than previously hypothesized by Titus et al. (2017b) and highlights a novel diversification pathway that has generated species-level diversity across a relatively small, but biodiverse, marine region.

**Figure 4.**
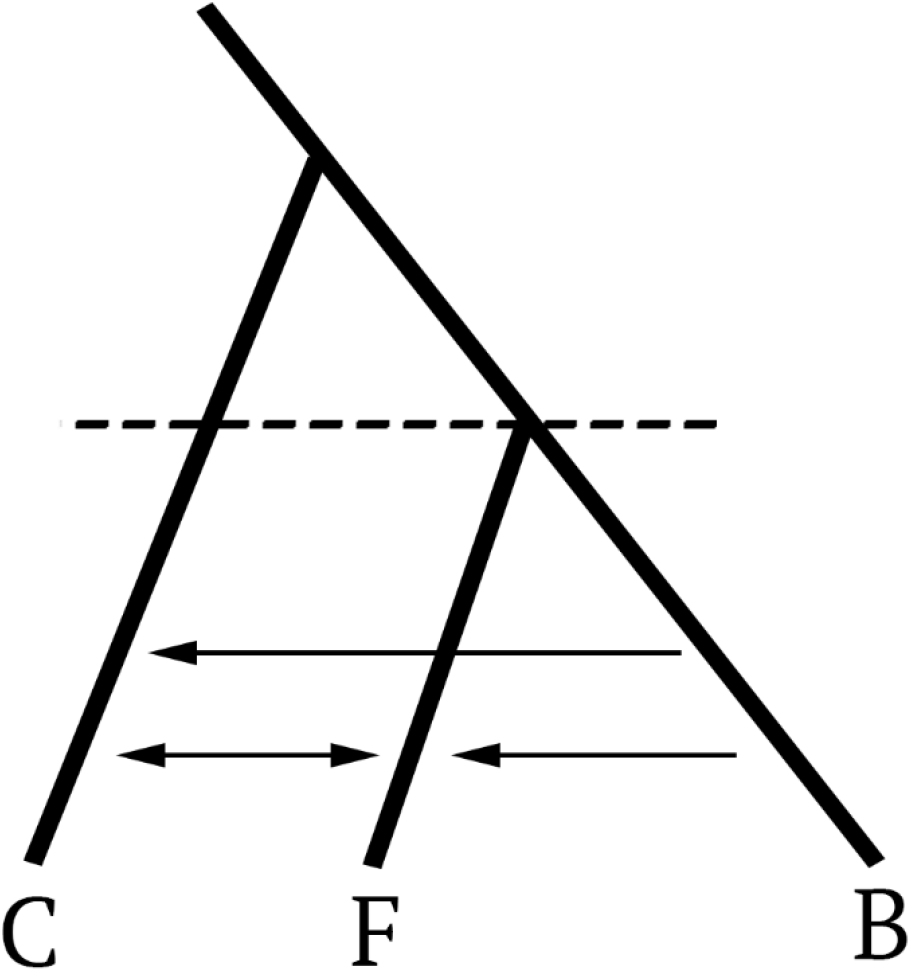
Best fit topology and evolutionary model for the *Ancylomenes pedersoni* species complex. Arrows represent migration events in the coalescent (i.e. backwards in time).

The formation of the IOP is one of the most globally significant and heavily studied, natural events in Earth’s history (reviewed by O’Dea et al. 2016). While the resulting land bridge facilitated terrestrial range expansions and faunal interchange between North and South America, in the oceans, the IOP severed the open seaway between the Atlantic and Pacific Oceans and sent their respective communities on independent evolutionary trajectories (Knowlton et al., 1993; Lessios 2008). In the Western Atlantic, the Caribbean Sea became warmer, saltier, and oligotrophic, thereby facilitating major coral reef development (Lessios 2008). Although a pulse of ecological speciation in scleractinian corals occurred in the 1 million years following the formation of the IOP, the IOP fundamentally changed ocean circulation and global climate patterns, leading directly to northern hemisphere glaciation and a period of marine extinction (Budd and Johnson 1999). The repeated reduction in shallow water habitat that resulted from the Pleistocence glacial cycles also decreased sea surface temperatures and drove major extinctions in Caribbean corals (Budd and Johnson 1999). The result led to major ecological turnover, and opportunity, in the Greater Caribbean (Budd and Johnson 1999; Budd and Klaus 2001; Prada et al. 2016). In fact, while the origins of many higher-level taxonomic groups in the Caribbean can be traced to major vicariant events such as the closure of the Tethys Sea and the formation of the IOP (Cowman et al. 2013; Bowen et al. 2016), much of the species-level diversity that has evolved within the Caribbean Sea itself appears to have originated ecologically/sympatrically rather than allopatrically. Examples are wide-ranging and include reef fishes (Ruber et al. 2003; Barber and Bellwood 2005; Rocha et al. 2005; Taylor and Hellberg 2005; Rocha et al. 2008; Tavera et al. 2012), scleractinian corals (Weil and Knowlton 1994; Carlon and Budd, 2007; Frade et al. 2010), octocorals (Lasker et al. 1983; Prada and Hellberg 2013; Prada et al. 2014; Prada and Hellberg 2020), crustaceans (Knowlton and Keller 1983, 1985; Hurt et al. 2013), and sea anemones (Titus et al. 2019a).

In contrast, allopatric speciation within the Greater Caribbean appears to be much rarer. The Mona Passage between Hispañola and Puerto Rico is identified most frequently as the major phylogeographic barrier in the region separating Western and Eastern Caribbean regions, but this break is almost exclusively resolved at the intraspecific level (reviewed by DeBiasse et al., 2016). Hyper-restricted endemic taxa have been only been described occasionally from the Meso-American Barrier Reef and Southern Caribbean (Colin, P. 2002; Taylor and Hellberg 2005; 2006; Hurt et al. 2013; D’Aloia et al. 2017). Much of the allopatric diversity that arises in the Tropical Western Atlantic comes via dispersal from the Greater Caribbean to Brazil where major freshwater runoff from the Amazon basic creates a major barrier to gene flow between regions (reviewed by Bowen et al. 2016). This is despite the fact that the formation of the IOP deflected equatorial currents in the Atlantic northwards through the Caribbean Sea and into the Gulf of Mexico where the current loops clockwise, mounds slightly, and then is pulled downwards by gravity through the narrow Florida Straits, strengthening the Gulf Stream, which can move 30 million cubic meters of water per second at velocities up to 2.5 m/s (Fig 1). Although a number of phylogeographic studies also identify the Florida Straits as an important intraspecific phylogeographic barrier, to our knowledge, no examples have demonstrated that speciation has occurred across this barrier, let alone coincided with the formation of the IOP and led to downstream endemic hybrid speciation. Instead, the Gulf Stream is often discussed as a means of explaining the relative homogeneity between Bermudan and Caribbean marine communities, and the subsequent low rates of Bermudan endemicity although though the nearest coral reef system is >1500 kilometers away (reviewed by Titus et al. 2017b).

Our data thus uncover a novel example of the evolutionary consequences the formation of the Isthmus of Panama has had on marine biodiversity. Our study is a particularly important example for marine evolutionary biologists demonstrating the complexity of the geological, biological, and physical oceanographic process that generate biodiversity on coral reefs reinforcing that ocean currents can act as temporal allopatric barriers and downstream dispersal pathways simultaneously. Finally, although introgressive hybridization is not uncommon in marine systems, true hybrid speciation is rarely documented, and the importance of these reticulate evolutionary processes are poorly understood (reviewed by Arnold and Fogarty 2009). The results of our study argue strongly that methods that model gene flow and reticulate processes be included regularly into systematic studies in the sea, particularly on the shallow evolutionary timescales we have worked with here.

## Acknowledgements

We thank Erich Bartels, Annelise del Rio, Jose Diaz, Dan Exton, Natalie Hamilton, Alex Hunter, Anna Klompen Jason Macrander, Spencer Palombit, Stephen Ratchford, Jill Titus, Cory Walter, Eric Witt, Clay Vondriska, and the Operation Wallacea dive staff for assistance in the field and laboratory. Alonso Delgado assisted with ddRADseq analyses, and Jordan Satler and Megan Smith provided valuable advice on simulation analyses and model selection. Bellairs Research Station, the Bermuda Institute of Ocean Science, Cape Eleuthera Institute, CARMABI, Coral View Dive Center, Gerace Research Centre, the Honduran Coral Reef Foundation, Mote Marine Laboratory, Smithsonian Tropical Research Institute, and the University of the Virgin Islands provided valuable logistical support. Specimens were collected under permits: SE/A-88-15, PPF/DGOPA-127/14, CZ01/9/9, FKNMS-2012-155, SAL-12-1432A-SR, STT037-14, 140408, MAR/FIS/17, 19985, and N. PPF/DGOPA-295/17. This research was supported by National Science Foundation Doctoral Dissertation Improvement Grant DEB 1601645 and Florida Fish and Wildlife Conservation Commission awards to B.M.T. & M.D. Operation Wallacea, American Philosophical Society, International Society for Reef Studies Graduate Fellowship, PADI Foundation Grant, American Museum of Natural History Lerner Gray Funds, and The Ohio State University Presidential Fellowship provided funding to B.M.T. Field work in Mexico was financially supported by CONACyTCB-2012-01-177293, CONABIO NE018, and the Harte Research Institute assigned to NS. Additional funding was provided through the Trautman Fund of The OSU Museum of Biological Diversity, The Ohio State University, and the National Science Foundation DEB-1257796 to MD.

## References

Álvarez-Noriega, M., Burgess, S.C., Byers, J.E., Pringle, J.M., Wares, J.P. and Marshall, D.J., 2020. Global biogeography of marine dispersal potential. Nature Ecology & Evolution, 4(9), pp.1196–1203.

Arnold, M.L. and Fogarty, N.D., 2009. Reticulate evolution and marine organisms: the final frontier? International Journal of Molecular Sciences, 10(9), pp.3836–3860.

Barber, P.H. and Bellwood, D.R., 2005. Biodiversity hotspots: evolutionary origins of biodiversity in wrasses (Halichoeres: Labridae) in the Indo-Pacific and new world tropics. Molecular Phylogenetics and Evolution, 35(1), pp.235–253.

Blischak, P.D., Chifman, J., Wolfe, A.D. and Kubatko, L.S., 2018. HyDe: a Python package for genome-scale hybridization detection. Systematic Biology, 67(5), pp.821–829.

Bowen, B.W., Rocha, L.A., Toonen, R.J. and Karl, S.A., 2013. The origins of tropical marine biodiversity. Trends in Ecology & Evolution, 28(6), pp.359–366.

Bowen, B.W., Gaither, M.R., DiBattista, J.D., Iacchei, M., Andrews, K.R., Grant, W.S., Toonen, R.J. and Briggs, J.C., 2016. Comparative phylogeography of the ocean planet. Proceedings of the National Academy of Sciences, 113(29), pp.7962–7969.

Briones-Fourzán, P., Pérez-Ortiz, M., Negrete-Soto, F., Barradas-Ortiz, C. and Lozano-Álvarez, E., 2012. Ecological traits of Caribbean sea anemones and symbiotic crustaceans. Marine Ecology Progress Series, 470, pp.55–68.

Bryant, D., Bouckaert, R., Felsenstein, J., Rosenberg, N.A. and RoyChoudhury, A., 2012. Inferring species trees directly from biallelic genetic markers: bypassing gene trees in a full coalescent analysis. Molecular Biology and Evolution, 29(8), pp.1917–1932.

Budd, A.F. and Johnson, K.G., 1999. Origination preceding extinction during late Cenozoic turnover of Caribbean reefs. Paleobiology, 25(2), pp.188–200.

Budd, A.F. and Klaus, J.S., 2001. The origin and early evolution of the Montastraea “annularis” species complex (Anthozoa: Scleractinia). Journal of Paleontology, 75(3), pp.527–545.

Burnham, K.P. and Anderson, D.R., 2002. Model selection and multimodal inference: A practical information theoretic approach. 2^nd^ ed. New York, NY: Springer

Carlon, D.B. and Budd, A.F., 2002. Incipient speciation across a depth gradient in a scleractinian coral? Evolution, 56(11), pp.2227–2242.

Chace, F.A. Jr. (1958) A new shrimp of the genus Periclimenes from the West Indies. Proceedings of the Biological Society of Washington, 71, 125–130.

Choat, J.H., 2006. Phylogeography and reef fishes: bringing ecology back into the argument. Journal of Biogeography, 33(6), pp.967–968.

Chifman, J. and Kubatko, L., 2014. Quartet inference from SNP data under the coalescent model. Bioinformatics, 30(23), pp.3317–3324.

Clement, M., Posada, D.C.K.A. and Crandall, K.A., 2000. TCS: a computer program to estimate gene genealogies. Molecular Ecology, 9(10), pp.1657–1659.

Colin, P.L., 2002. A new species of sponge-dwelling Elacatinus (Pisces: Gobiidae) from the western Caribbean. Zootaxa, 106(1), pp.1–7.

Cowman, P.F. and Bellwood, D.R., 2013. Vicariance across major marine biogeographic barriers: temporal concordance and the relative intensity of hard versus soft barriers. Proceedings of the Royal Society B: Biological Sciences, 280(1768), p.20131541.

D’Aloia, C.C., Bogdanowicz, S.M., Harrison, R.G. and Buston, P.M., 2017. Cryptic genetic diversity and spatial patterns of admixture within Belizean marine reserves. Conservation Genetics, 18(1), pp.211–223.

DeBiasse, M.B., Richards, V.P., Shivji, M.S. and Hellberg, M.E., 2016. Shared phylogeographical breaks in a Caribbean coral reef sponge and its invertebrate commensals. Journal of Biogeography, 43(11), pp.2136–2146.

Degnan, J.H. and Rosenberg, N.A., 2009. Gene tree discordance, phylogenetic inference and the multispecies coalescent. Trends in Ecology & Evolution, 24(6), pp.332–340.

Drummond, A.J. and Rambaut, A., 2007. BEAST: Bayesian evolutionary analysis by sampling trees. BMC Evolutionary Biology, 7(1), pp.1–8.

Eaton, D.A., 2014. PyRAD: assembly of de novo RADseq loci for phylogenetic analyses. Bioinformatics, 30(13), pp.1844–1849.

Edgar, R.C., 2004. MUSCLE: multiple sequence alignment with high accuracy and high throughput. Nucleic Acids Research, 32(5), pp.1792–1797.

Excoffier, L., Dupanloup, I., Huerta-Sánchez, E., Sousa, V.C. and Foll, M., 2013. Robust demographic inference from genomic and SNP data. PLoS Genet, 9(10), p.e1003905.

Frade, P.R., Reyes-Nivia, M.C., Faria, J., Kaandorp, J.A., Luttikhuizen, P.C. and Bak, R.P.M., 2010. Semi-permeable species boundaries in the coral genus Madracis: introgression in a brooding coral system. Molecular Phylogenetics and Evolution, 57(3), pp.1072–1090.

Haug, G.H. and Tiedemann, R., 1998. Effect of the formation of the Isthmus of Panama on Atlantic Ocean thermohaline circulation. Nature, 393(6686), pp.673–676.

Huebner, L.K., Shea, C.P., Schueller, P.M., Terrell, A.D., Ratchford, S.G. and Chadwick, N.E., 2019. Crustacean symbiosis with Caribbean sea anemone *Bartholomea annulata*: occupancy modeling, habitat partitioning, and persistence. Marine Ecology Progress Series, 631, pp.99–116.

Hurt, C., Silliman, K., Anker, A. and Knowlton, N., 2013. Ecological speciation in anemone-associated snapping shrimps *(Alpheus armatus* species complex). Molecular Ecology, 22(17), pp.4532–4548.

Gilpin, J.A. and Chadwick, N.E., 2017. Life-history traits and population structure of Pederson cleaner shrimps *Ancylomene spedersoni*. The Biological Bulletin, 233(3), pp.190–205.

Jiao, X., Flouri, T., Rannala, B. and Yang, Z., 2020. The impact of cross-species gene flow on species tree estimation. Systematic Biology, 69(5), pp.830–847.

Jombart, T., Devillard, S. and Balloux, F., 2010. Discriminant analysis of principal components: a new method for the analysis of genetically structured populations. BMC Genetics, 11(1), p.94.

Jombart, T. and Ahmed, I., 2011. adegenet 1.3-1: new tools for the analysis of genome-wide SNP data. Bioinformatics, 27(21), pp.3070–3071.

Kearse, M., Moir, R., Wilson, A., Stones-Havas, S., Cheung, M., Sturrock, S., Buxton, S., Cooper, A., Markowitz, S., Duran, C. and Thierer, T., 2012. Geneious Basic: an integrated and extendable desktop software platform for the organization and analysis of sequence data. Bioinformatics, 28(12), pp.1647–1649.

Keith, N., Tucker, A.E., Jackson, C.E., Sung, W., Lledó, J.I.L., Schrider, D.R., Schaack, S., Dudycha, J.L., Ackerman, M., Younge, A.J. and Shaw, J.R., 2016. High mutational rates of large-scale duplication and deletion in *Daphnia pulex*. Genome Research, 26(1), pp.60–69.

Knowlton, N. and Keller, B.D., 1983. A new, sibling species of snapping shrimp associated with the Caribbean sea anemone *Bartholomea annulata*. Bulletin of marine science, 33(2), pp.353–362.

Knowlton, N. and Keller, B.D., 1985. Two more sibling species of alpheid shrimps associated with the Caribbean sea anemones *Bartholomea annulata* and *Heteractis lucida*. Bulletin of Marine Science, 37(3), pp.893–904.

Knowlton, N., Weigt, L.A., Solorzano, L.A., Mills, D.K. and Bermingham, E., 1993. Divergence in proteins, mitochondrial DNA, and reproductive compatibility across the Isthmus of Panama. Science, 260(5114), pp.1629–1632.

Lasker, H.R., Gottfried, M.D. and Coffroth, M.A., 1983. Effects of depth on the feeding capabilities of two octocorals. Marine Biology, 73(1), pp.73–78.

Leaché, A.D., Fujita, M.K., Minin, V.N. and Bouckaert, R.R., 2014. Species delimitation using genome-wide SNP data. Systematic Biology, 63(4), pp.534–542.

Leaché, A.D., Harris, R.B., Rannala, B. and Yang, Z., 2014. The influence of gene flow on species tree estimation: a simulation study. Systematic Biology, 63(1), pp.17–30.

Lessios, H.A., 2008. The great American schism: divergence of marine organisms after the rise of the Central American Isthmus. Annual Review of Ecology, Evolution, and Systematics, 39, pp.63–91.

Mascaró, M., Rodríguez-Pestaña, L., Chiappa-Carrara, X. and Simões, N., 2012. Host selection by the cleaner shrimp *Ancylomenes pedersoni:* Do anemone host species, prior experience or the presence of conspecific shrimp matter? Journal of Experimental Marine Biology and Ecology, 413, pp.87–93.

O’Dea, A., Lessios, H.A., Coates, A.G., Eytan, R.I., Restrepo-Moreno, S.A., Cione, A.L., Collins, L.S., De Queiroz, A., Farris, D.W., Norris, R.D. and Stallard, R.F., 2016. Formation of the Isthmus of Panama. Science Advances, 2(8), p.e1600883.

Palumbi, S.R., 1994. Genetic divergence, reproductive isolation, and marine speciation. Annual Review of Ecology, Evolution, and Systematics, 25(1), pp. 547–572.

Potkamp, G. and Fransen, C.H., 2019. Speciation with gene flow in marine systems. Contributions to Zoology, 88(2), pp.133–172.

Prada, C. and Hellberg, M.E., 2013. Long prereproductive selection and divergence by depth in a Caribbean candelabrum coral. Proceedings of the National Academy of Sciences, 110(10), pp.3961–3966.

Prada, C., McIlroy, S.E., Beltrán, D.M., Valint, D.J., Ford, S.A., Hellberg, M.E. and Coffroth, M.A., 2014. Cryptic diversity hides host and habitat specialization in a gorgonian algal symbiosis. Molecular Ecology, 23(13), pp.3330–3340.

Prada, C., Hanna, B., Budd, A.F., Woodley, C.M., Schmutz, J., Grimwood, J., Iglesias-Prieto, R., Pandolfi, J.M., Levitan, D., Johnson, K.G. and Knowlton, N., 2016. Empty niches after extinctions increase population sizes of modern corals. Current Biology, 26(23), pp.3190–3194.

Prada, C. and Hellberg, M., 2020. Speciation-by-depth on coral reefs: sympatric divergence with gene flow or cryptic transient isolation? Journal of Evolutionary Biology.

Puillandre, N., Lambert, A., Brouillet, S. and Achaz, G., 2012. ABGD, Automatic Barcode Gap Discovery for primary species delimitation. Molecular Ecology, 21(8), pp.1864–1877.

Quenouille, B., Hubert, N., Bermingham, E. and Planes, S., 2011. Speciation in tropical seas: allopatry followed by range change. Molecular Phylogenetics and Evolution, 58(3), pp.546–552.

R Core Team 2015. R: A language and environment for statistical computing. Vienna, Austria: R Foundation for Statistical Computing. https://www.R-project.org/

Rannala, B., Edwards, S.V., Leaché, A. and Yang, Z., 2020. The multi-species coalescent model and species tree inference. Phylogenetics in the Genomic Era, pp.3.3:1-3.3:21.

Rocha, L.A., Robertson, D.R., Roman, J. and Bowen, B.W., 2005. Ecological speciation in tropical reef fishes. Proceedings of the Royal society B: Biological Sciences, 272(1563), pp.573–579.

Rüber, L., Van Tassell, J.L. and Zardoya, R., 2003. Rapid speciation and ecological divergence in the American seven-spined gobies (Gobiidae, Gobiosomatini) inferred from a molecular phylogeny. Evolution, 57(7), pp.1584–1598.

Simmonds, S.E., Chou, V., Cheng, S.H., Rachmawati, R., Calumpong, H.P., Mahardika, G.N. and Barber, P.H., 2018. Evidence of host-associated divergence from coral-eating snails (genus *Coralliophila)* in the Coral Triangle. Coral Reefs, 37(2), pp.355–371.

Smith, M.L., Ruffley, M., Espíndola, A., Tank, D.C., Sullivan, J. and Carstens, B.C., 2017. Demographic model selection using random forests and the site frequency spectrum. Molecular Ecology, 26(17), pp.4562–4573.

Solís-Lemus, C., Yang, M. and Ané, C., 2016. Inconsistency of species tree methods under gene flow. Systematic Biology, 65(5), pp.843–851.

Stamatakis, A., 2014. RAxML version 8: a tool for phylogenetic analysis and post-analysis of large phylogenies. Bioinformatics, 30(9), pp.1312–1313.

Swofford, D.L., 2004. Paup 4.0 for Macintosh: Phylogenetic analysis using parsimony (software and user’s book for Macintosh)

Tavera, J.J., Acero, A., Balart, E.F. and Bernardi, G., 2012. Molecular phylogeny of grunts (Teleostei, Haemulidae), with an emphasis on the ecology, evolution, and speciation history of New World species. BMC Eolutionary Biology, 12(1), p.57.

Taylor, M.S. and Hellberg, M.E., 2005. Marine radiations at small geographic scales: speciation in neotropical reef gobies *(Elacatinus)*. Evolution, 59(2), pp.374–385.

Taylor, M.S. and Hellberg, M.E., 2006. Comparative phylogeography in a genus of coral reef fishes: biogeographic and genetic concordance in the Caribbean. Molecular Ecology, 15(3), pp.695–707.

Titus, B.M. and Daly, M., 2015. Fine-scale phylogeography reveals cryptic biodiversity in Pederson’s cleaner shrimp, *Ancylomenes pedersoni* (Crustacea: Caridea: Palaemonidae), along the Florida Reef Tract. Marine Ecology, 36(4), pp.1379–1390.

Titus, B.M. and Daly, M., 2017. Specialist and generalist symbionts show counterintuitive levels of genetic diversity and discordant demographic histories along the Florida Reef Tract. Coral Reefs, 36(1), pp.339–354.

Titus, B.M., Daly, M. and Exton, D.A., 2015. Temporal patterns of Pederson shrimp *(Ancylomenes pedersoni* Chace 1958) cleaning interactions on Caribbean coral reefs. Marine Biology, 162(8), pp.1651–1664.

Titus, B.M., Vondriska, C. and Daly, M., 2017a. Comparative behavioural observations demonstrate the ‘cleaner’shrimp *Periclimenes yucatanicus* engages in true symbiotic cleaning interactions. Royal Society Open Science, 4(4), p.170078.

Titus, B.M., Palombit, S. and Daly, M., 2017b. Endemic diversification in an isolated archipelago with few endemics: an example from a cleaner shrimp species complex in the Tropical Western Atlantic. Biological Journal of the Linnean Society, 122(1), pp.98–112.

Titus, B.M., Blischak, P.D. and Daly, M., 2019a. Genomic signatures of sympatric speciation with historical and contemporary gene flow in a tropical anthozoan (Hexacorallia: Actiniaria). Molecular Ecology, 28(15), pp.3572–3586.

Titus, B.M., Daly, M., Vondriska, C., Hamilton, I. and Exton, D.A., 2019b. Lack of strategic service provisioning by Pederson’s cleaner shrimp *(Ancylomenes pedersoni)* highlights independent evolution of cleaning behaviors between ocean basins. Scientific Reports, 9(1), pp.1–9.

Vermeij, G.J. and Grosberg, R.K., 2010. The great divergence: when did diversity on land exceed that in the sea? Integrative and Comparative Biology, 50(4), pp.675–682.

Weil, E. and Knowton, N., 1994. A multi-character analysis of the Caribbean coral *Montastraea annularis* (Ellis and Solander, 1786) and its two sibling species, *M. faveolata* (Ellis and Solander, 1786) and *M. franksi* (Gregory, 1895). Bulletin of Marine Science, 55(1), pp.151–175.

