## Supplementary Material for "Topology Testing and Demographic Modeling Illuminate a Novel Speciation Pathway in the Greater Caribbean Sea Following the Formation of the Isthmus of Panama"

**­­­**

**Supplementary Methods**

*Sampling and DNA Extraction*

Whole animals of *Ancylomenes pedersoni* were collected from sea anemones, primarily *Bartholomea annulata*, from 14 coral reef sites representing the entire geographic range of the species complex. Individuals were collected by hand, using SCUBA, between 3-20 meters depth using Whirl-Pak sample bags. All individuals were preserved on shore using 95% EtOH or RNAlater. Samples from the Florida Keys and Bermuda had been collected previously (N = 131 individuals; see Titus and Daly 2015, 2017; Titus et al. 2017b). All other individuals were newly collected for this study (N = 311 individuals). Additionally, two individuals of *Periclimenes yucatanicus*, a closely related species, were collected from Curacao and included as outgroup taxa for genomic analyses.

All samples were transported back to The Ohio State University where total genomic DNA was extracted from abdominal tissue using QIAGEN DNeasy Blood & Tissue Kits and stored at -20°C. DNA extractions were vacuum centrifuged to 50uL total volume and then quantified (ng/μL) using Qubit 2.0 (ThermoFisher) fluorometer and dsDNA broad-range assay kits.

*Sequencing and Dataset Assembly*

For each *A. pedersoni* individual we PCR amplified a 650bp long DNA fragment from cytochrome *c* oxidase subunit I (COI) using the universal primers LCO1490 and HCO2198. PCR run conditions can be found in Titus and Daly (2015). mtDNA barcode sequence data were assembled using Sequencher 4.9 (Gene Codes) and aligned using MUSCLE (Edgar 2004) in Geneious v10.2.3 (Kearse et al. 2012). The crustacean origin of each sequence was verified using nucleotide BLAST query in GenBank. Following alignment, nucleotide sequences were translated into amino acids to look for stop codons within the open reading frame as nuclear COI pseudogenes are commonly found in crustaceans. Previously published COI sequence data from five shrimp species in the genus *Periclimenes* were included as outgroup taxa: *P. yucatanicus* (KX926215- KX926217; Titus and Daly 2017), *P. rathbunae* (KX090125, KX090114, KU065005; Brinkman and Fransen 2016; Horka et al., 2016); *P. crinoidalis* (KU064995; Horka et al., 2016), *P. perryae* (KU065004; Horka et al., 2016), and *P. patae* (KU065002; Horka et al. 2016).

Following DNA extraction and quantification, DNA degradation was assessed for each individual sample using gel electrophoresis. For each sample locality, 10-15 samples with high molecular weight DNA were carried forward for ddRADseq library preparation. We followed the ddRADseq preparation protocol of Sovic et al., (2016). For each sample 20μL aliquots, each with 200ng of DNA were digested using the restriction enzymes *Eco-*RI-HF and *PstI*-HF enzymes. Illumina compatible barcodes were annealed to the overhanging ends of each DNA fragment. Samples were size selected manually using a 400-800 bp size range, and then cleaned using Nucleospin Gel and PCR clean up kits (Machery-Nagel). Following size selection, each sample was amplified using PCR, cleaned using AMpure XP beads (Agilent), and quantified using qPCR to inform the pooling of individual samples into final libraries. A total of 152 *A. pedersoni* individuals passed all quality control steps and were pooled into 8 separate libraries and sequenced across 8 separate Illumina lanes. Sequencing was conducted using 100bp single-end sequencing on an Illumina HiSeq2500 at the James Comprehensive Cancer Center Genomics Shared Resource at The Ohio State University.

Raw ddRADseq reads were demultiplex, aligned, and assembled *de novo* using the program pyRAD v 3.066 (Eaton 2014). We set the maximum number of low-quality base calls (max_low_qual_bases) in a read to five, and low quality base calls (Q < 20) were replaced with Ns. The Phred Q score offset (phred_Qscore_offset) was set to 33. We set the clustering threshold (Wclust) to 0.90 to assemble reads into loci. We did not allow any mismatches in barcode sequences, and elected to strictly filter adapters (filter = 2 parameter setting). The maximum number of alleles in a consensus sequence was set to 2, and the maximum number of heterozygous sites in a consensus sequence was set to 5 (i.e. 5% of all sites in a 100bp sequence) in order to remove poor alignments. We allowed up to 20% SNPs per locus (maxSNPs = 0.2) and set the maximum number of indels per locus to 8 (max_Indels_locus). The maximum number of individuals with a shared heterozygous site was set to 50%. We required a locus to be present in 75% of all individuals to be retained in the final dataset.

Because different downstream analyses may or may not require outgroups, and because individual variation in sequencing coverage can influence the number of recovered SNPs, we created three ddRADseq datasets. Our first dataset included all *A. pedersoni* individuals that had >750,000 raw Illumina sequence reads and at least 10,000 consensus loci after clustering. (N = 135 individuals; Table S2). This comprehensive range-wide dataset was assembled, like our mtDNA dataset, to confirm we had captured all the species-level diversity in the *A. pedersoni* species complex throughout the entire Tropical Western Atlantic. However, this dataset traded decreased SNP recovery for increased sample sizes (N = 1232 SNPs and 135 individuals; Table S2) which was a tradeoff we felt was important to make at the “species discovery” stage. After confirming we had captured all species level diversity, the second dataset we assembled was a reduced *A. pedersoni* dataset that included two individuals of *Periclimenes yucatanicus* as outgroup taxa. This dataset was built to create even sample sizes across each of the three *A. pedersoni* lineages for conducting phylogenetic analyses (N = 15 from Bermuda and Caribbean lineages; N = 13 from Floridian lineage). In total our reduced dataset with outgroup taxa included N = 45 individuals and 2101 SNPs (Table S2). Finally, our third dataset includes the same individuals as the reduced dataset but without including outgroup taxa. This dataset was built to increase the number of recovered SNPs across each of the three *A. pedersoni* lineages in order to perform demographic modeling using coalescent simulations and the multidimensional site frequency spectrum (mSFS). In total our reduced dataset without outgroup taxa included N = 43 individuals and N = 4053 SNPs (Table S2).

*Species Delimitation and Phylogenetic Analyses*

To confirm that Titus et al (2017) had identified all *A. pedersoni* species-level diversity throughout the Tropical Western Atlantic we conducted species delimitation analyses for both single locus and ddRADseq datasets. Using COI DNA barcodes, we used two species delimitation approaches: 1) pairwise sequence divergence using the program Automatic Barcode Gap Discovery (ABGD; Puillandre et al. 2012), and 2) statistical parsimony networks using the program TCS v2.1 (Clement et al. 2000). Neither program uses *a priori* species assignments to guided delimitations, and thus, each presents a broad snapshot of the genetic divergence within a sample of sequences that can then be more rigorously tested to determine whether distinct genetic groups are divergent enough to warrant status as separate species. A 3% pairwise sequence divergence estimate was used as a threshold in ABGD to distinguish putative intraspecific and interspecific divergence following common practice for crustaceans (Plaisaince et al., 2009). In TCS we used a 95% parsimoniously plausible branch connections to distinguish putative species. Cryptic diversity within a sample of sequences that exceed this threshold form unconnected networks and have been found to represent true species level divergence (Hart and Sunday 2007).

With our ddRADseq dataset we searched for species-level genetic partitions in our data using genetic cluster analyses (Carstens et al. 2013). We used discriminate analyses of principle components (DAPC; Jombart et al. 2010) in the adegenet package (Jombart and Ahmed 2011) in R v3.5.0 (R Core Team 2015). The optimal number of genetic clusters was determined using the *K*-means method, setting the maximum *K* = 10, and retaining all principal components. The most likely number of *K* genetic clusters was defined using the lowest Bayesian Information Criteria (BIC) value. The genetic data were then transformed using principal component analysis (PCA) and a linear discriminant analysis was performed on the retained principal components. No more than 50% of the PCs were retained to avoid overfitting. We then assigned each individual toa genetic unit according to its maximum membership probability.

Both single-locus and ddRADseq data recovered three deep genetic partitions, and a fourth shallow genetic break within the Caribbean lineage that corresponds to a commonly recovered intraspecific phylogeographic break at the Mona Passage (see Results). Thus, we focused our remaining species delimitation analyses using the three deepest species level partitions in our data. Using SNP data we conducted species delimitation analyses with path sampling and Bayes factors in the program SNAPP and Bayes Factor Delimitation* (BFD*; Leache et al. 2014). BFD* uses unlinked SNPs and marginal likelihood estimated calculated via path sampling to perform model selection on competing species delimitation models (Bryant et al. 2012). Specifically, we test the current taxonomy of *A. pedersoni* (i.e. one widespread species) versus species delimitations recovered by single and multi-locus species discovery analyses (i.e. two and three species models, respectively). Due to the computational constraints of running SNAPP with biallelc data (i.e. computation time increases linearly as more loci are added but exponentially as more samples are added), and constraints for estimating species trees with missing data (i.e. data must be present in at least one individual at each locus for each putative species, we conducted our analyses with only 11 individuals so that analyses could be completed over the course of days rather than weeks or months. We also used three individuals from each putative *A. pedersoni* lineage plus two *P. yucatanicus* individuals as outgroups (Table S3). In pyRAD, we required a locus to be present in 10 of 11 individuals to meet the assumptions of SNAPP. For each SNAPP analysis, mutation rates *u* and *v* were set to 1 and were not sampled, and the coalescent rate was set at 10 and sampled throughout the analysis. We used only polymorphic loci and a broad gamma distributed prior (2,200) for speciation rate (λ). We conducted the path analysis (48 steps) in SNAPP v1.3 and BEAST v 2.4.6 for 1 x 10^5^ MCMC generations with 10% discarded as burnin (Bouckaert et al. 2014). MCMC chains were assessed using Tracer v1.6.0 and considered properly mixed when the effective sample size for all parameters > 200 (Heled and Drummond 2010). All Bayes factor calculations were made against the current model of *A. pedersoni* as a single widespread species in the Tropical Western Atlantic. Positive Bayes factors indicate support for the current model, while negative Bayes factors indicate support for the alternative model. Where multiple species delimitation models outperform the current model, the model with the lowest Bayes factor is considered the best fit to the data. Finally, We used Arlequin v 3.5 (Excoffier and Lisher 2010) to compute measures of genetic differentiation (F_ST_) between putative lineages.

*Topology testing and demographic model selection*

Historical and contemporary gene flow can affect species tree topologies and parameter estimations. However, species tree analyses use the multi-species coalescent model, which assumes gene tree discordance is the result of incomplete lineage sorting (ILS) rather than gene flow. Our mitochondrial gene trees and ddRADseq species trees produced conflicting, yet fully supported, tree topologies, and genetic clustering analyses suggest ongoing introgression where Floridian and Caribbean *A. pedersoni* lineages co-occur along the Florida Reef Tract (see Results). Thus, historical and contemporary gene flow is likely important to account for when reconstructing the evolutionary history of this group. To do this, we incorporated migration parameters into topology testing and demographic model selection analyses to provide statistically supported inference on the evolutionary scenarios that may have given rise to the *A. pedersoni* species complex. We used Dataset 3 (43 individuals with no outgroup; Table S2), the multidimensional site frequency spectrum (mSFS) and coalescent simulations in *fastsimcoal2* (FSC2; Excoffier et al. 2013) for this purpose. We built a comprehensive set of 45 evolutionary models across all possible species tree topologies that vary in the timing and directionality of migration between putative species.

Twelve models were built for each possible species tree topology: ((Caribbean, (Florida, Bermuda)), ((Florida, (Caribbean, Bermuda)), and ((Bermuda, (Florida, Caribbean)). Models tested for contemporary and ancestral gene flow between all putative species, but given Bermuda’s geographic location and directionality of the Gulf Stream, we only tested one model with gene flow from Bermuda to the greater Caribbean for each tree topology. All other models specified unidirectional gene flow to Bermuda. We also tested an additional nine evolutionary scenarios that are not explicitly reliant on hierarchical species tree topologies: three island models testing all migration combinations, three models with and without migration where the Caribbean, Florida, and Bermudan species diverged simultaneously forming an unresolved polytomy, and three hybrid speciation models, with and without migration, where both the Caribbean and Florida species provided 50% of the genetic material that gave rise to the endemic Bermudan species (Fig. 3).

Using output from pyRAD and custom python scripts from Satler et al. (2017) and Smith et al. (2017), we constructed a 3-population joint-folded mSFS for putative *A. pedersoni* lineages from Florida, Caribbean, and Bermuda. mSFS calculations require fixed numbers of alleles from all populations (i.e. no missing data). Meeting this requirement would have greatly decreased our dataset size and likely biased our analyses. To deal with missing data, yet incorporate the greatest number of SNPs into the mSFS as possible, we employed a down sampling approach with a threshold of 75% following Satler et al. (2017) and Smith et al. (2017). This approach only uses SNPs that are present in at least 75% of all individuals to create the mSFS. Loci that exceed this threshold are randomly subsampled. Briefly, we built our mSFS as follows: 1) if a locus had fewer alleles than our threshold it was discarded, 2) if a locus had the exact number of alleles as the threshold, the minor allele frequency was recorded. 3) if a locus exceeded the threshold, alleles were downsampled with replacement until the number of alleles met the threshold, at which point the minor allele frequency was counted. This approach allowed us to maximize the number of SNPs included in the mSFS, but also has the potential to lead to monomorphic sites based on the downsampling procedure. To account for variation and possible monomorphic sites that could be the result of this down sampling procedure, we repeated the mSFS building procedure 10 times.

We used a mutation rate of 3.9e^-9^ substitutions per site per generation, calculated from the genome of the crustacean species *Daphnia pulex* (Order Cladocera; Keith et al. 2016), to convert parameter estimates to real values. Parameter estimates were then further scaled by assuming 2 generations per year because *A. pedersoni* reaches sexual maturity at ~6 months (Gilipin & Chadwick 2017). All models were repeated 50 times for each mSFS replicate (45 models x 10 mSFS replicates x 50 simulations/model/mSFS replicate = 22,500 simulations). The run with the highest composite likelihood, from each of the 50 runs, was selected for parameter estimation and model selection. The model with the best fit to the data was selected using AIC. Parameter estimates and 95% confidence intervals were then calculated for the best fit model.

*Tests of introgression, incomplete lineage sorting, and hybrid speciation*

Genetic cluster plots and model selection analyses suggest ongoing introgression between Floridian and Caribbean *A. pedersoni* lineages. Further, model selection results point to the importance of gene flow from both Floridian and Caribbean lineages in giving rise to the Bermudan endemic. Thus, we conducted ABBA/BABA and hybrid detection tests to independently confirm these inferences.

To disentangle introgression from ILS along the Florida Reef Tract, we conducted ABBA/BABA tests by calculating *D-*statistics after Patterson et al. (2012). The ABBA/BABA test assumes that SNP patterns of “ABBA” and “BABA” should be equally present between individuals from two populations that are diverging without gene flow, and thus, shared alleles are the result of ILS. Conversely, an excess of either pattern between individuals from two populations suggests introgression. ABBA/BABA tests were conducted in pyRAD with 200 bootstrap replicates with the following species tree recovered by SNAPP, SVDquartets, and FSC2 model selection: ((Bermuda, Florida), Caribbean). Observed *D-*statistics were converted to Z-scores, representing the number of standard deviations, from 0, the *D-*statistics deviated. As the ABBA/BABA test occurs between individuals rather than populations, we made multiple comparisons and used a *p*-value = 0.001 (Z-score > 3) to assess significance after correcting for multiple testing.

Finally, FSC2 model selection results highlight the importance of gene flow from both Floridian and Caribbean lineages in giving rise to the endemic Bermudan lineage. Thus, the Bermudan lineage may represent a hybrid species, or at minimum, the Caribbean lineage may have contributed a substantial portion of the observed Bermudan diversity (supported by the mtDNA gene tree as well showing the Bermudan mtDNA is derived from the Caribbean mtDNA). We independently test this hypothesis using the hybrid detection program HyDe (Blishak et al. 2017). HyDe uses phylogenetic invariants to implement a test for hybridization that is similar to the ABBA/BABA test, but can detect hybridization at both the individual and population level, estimate the proportion of the hybrid genome that came from each parental species, and test all possible hybridization scenarios. It thus allows us to test whether the Caribbean lineage contributed a substantial proportion of its genome, while the ABBA/BABA could not.

Table S1. Sample localities, GPS coordinates (Latitude and Longitude), *N*= sample sizes sequenced for COI barcodes and (ddRADseq), and GenBank Accession numbers for COI barcodes

| **Locality** | **Latitude** | **Longitude** | ***N*** | **GenBank Accession #s** |
| --- | --- | --- | --- | --- |
| Bocas del Toro, Panama | 9°19'45.29"N | 82°15'17.26"W | 27 (11) |  |
| Cayos Cochino, Honduras | 15°57'1.12"N | 86°29'51.82"W | 48 (6) |  |
| Utila, Honduras | 16° 5'18.03"N | 86°54'38.54"W | 37 (10) |  |
| Mahahual, Mexico | 18°42'18.45"N | 87°42'34.46"W | 49 (10) |  |
| Ft. Lauderdale, Florida | 26° 4'19.80"N | 80° 5'46.68"W | 15 (11) | KM064530-KM064544 |
| Upper Keys, Florida | 24°59'11.43"N | 80°24'55.83"W | 30 (11) | KM064592-KM064621 |
| Middle Keys, Florida | 24°41'58.09"N | 80°56'21.48"W | 8 (6) | KM064529-KM064530  KM064586-KM064591 |
| Lower Keys, Florida | 24°33'42.39"N | 81°23'31.59"W | 41 (19) | KM064545-KM064585 |
| Cape Eleuthera, Bahamas | 24°49'5.28"N | 76°20'51.80"W | 23 (12) |  |
| San Salvador, Bahamas |  |  | 9 (10) |  |
| St. Thomas, US Virgin Islands | 18°19'0.69"N | 64°59'22.59"W | 41 (10) |  |
| Barbados | 13°11'30.52"N | 59°38'29.04"W | 46 (10) |  |
| Curacao | 12° 7'19.45"N | 68°58'10.80"W | 40 (10) |  |
| Bermuda | 32°26'53.62"N | 64°45'45.42"W | 37 (15) | KZ845476-KX857630 |
| **Total** |  |  | **451 (151)** |  |

Table S2. Double Digest Restriction Site Associated DNA sequencing (ddRADseq) sequencing and assembly statistics. CSV file available on Dryad.

Table S3. *Ancylomenes pedersoni* samples used in Bayes Factor Delimitation* analyses.

| **Sample ID** | **Lineage** | **Loci** |
| --- | --- | --- |
| 2_LKAP22 | Florida | 1194 |
| 2_LKAP27 | Florida | 1181 |
| 2_LKAP41 | Florida | 1184 |
| FTAP4 | Caribbean | 1080 |
| UKAP59 | Caribbean | 1119 |
| UKAP6 | Caribbean | 1165 |
| BDAP14 | Bermuda | 1172 |
| BDAP15 | Bermuda | 1156 |
| BDAP34 | Bermuda | 1181 |
| CUPY11 | *Periclimenes yucatanicus* | 1045 |
| CUPY21 | *Periclimenes yucatanicus* | 1161 |
| **Total** |  | 1211 |

Table S4. Path sampling results for three species delimitation models for *Ancylomenes pedersoni* in the Tropical Western Atlantic. All Bayes Factor calculations are made against the model provided by current taxonomy. Positive Bayes Factors indicate support for this current model, whereas negative Bayes Factors indicate support for an alternative model. For each model *Periclimenes yucatanicus* was included as an outgroup. The highest ranked model is highlighted.

| **Model** | **Species** | **MLE** | **Rank** | **BF** |
| --- | --- | --- | --- | --- |
| *P. yucatancus* + current taxonomy | 2 | -8729.98 | 3 | - |
| *P. yucatanicus* + Bermuda + Caribbean | 3 | -8399.25 | 2 | -661.46 |
| *P. yucatanicus* + Bermuda + Caribbean + Florida | 4 | -8144.93 | 1 | -1170.1 |

MLE = Marginal Likelihood Estimate; BF = Bayes Factor

Table S5. Parameter estimates from FSC2 simulations for the *Ancylomenes pedersoni* species complex, calculated by the best fit model as selected by AIC in Table 8. Parameter estimates are scaled by a mutation rate of 3.9e^-9^ and 2 generations per year. FL = Florida, BD = Bermuda, CB = Caribbean. TDIV-FLBD = Divergence time between FL and BD. TDIV-FLBDCB = Divergence time between FL, BD, and CB. Arrows represent migration rates forward in time. Divergence time estimates are presented in years before present. Migration rates represent the effective number of migrants per year. CI = confidence interval

|  | **Effective population size (N_e_)** | | **Divergence time (**$\boldsymbol{\tau}$**)** | | **Migration** | |
| --- | --- | --- | --- | --- | --- | --- |
|  | Mean | 95% CI | Mean | 95% CI | Mean | 95% CI |
| Florida | 1,913,727 | ± 237,515 |  |  |  |  |
| Bermuda | 710 | ± 270 | - | - | - | - |
| Caribbean | 127,024 | ± 24,424 | - | - | - | - |
| FL+BD Ancestor | 232,767 | ± 66,921 | - | - | - | - |
| FL+BD+CB Ancestor | 1,490,871 | ± 471,290 | - | - | - | - |
| TDIV-FLBD | - | - | 202,968 | ± 21,522 | - | - |
| TDIV-FLBDCB | - | - | 3,528,268 | ± 503,770 | - | - |
| FL🡪 BD | - | - | - | - | 7.0e^-7^ | ± 1.8e^-6^ |
| FL 🡪 CB | - | - | - | - | 4.1e^-7^ | ± 4.7e^-8^ |
| CB🡪 BD | - | - | - | - | 7.9e^-7^ | ± 2.4e^-5^ |
| CB🡪 FL | - | - | - | - | 4.1e^-7^ | ± 3.7e^-8^ |

Table S6. Results from ABBA/BABA tests for *Ancylomenes pedersoni*. Only individuals with significant D-test results are reported. Individuals in **bold** are representatives indicated by red arrows in Figure 10. ABBA/BABA uses the following species tree topology ((P1, P2), P3) to test for introgression between species P2 and P3. Below, P1 = Bermuda, P2 = Florida, and P3 = Caribbean putative species. Out = outgroup represented by *Periclimenes yucatanicus*. D = Patterson’s D statistic. Z = results of two-tailed Z-test, testing for significantly positive D values. BABA and ABBA represent patterns of single nucleotide polymorphisms (SNPs) for each individual test of introgression (P1-P3). SNP patterns are assumed to be equally present between individuals diverging without gene flow, signifying shared alleles are the result of incomplete lineage sorting. Deviations from this expectation indicate introgression between P2 and P3.

| P1 | P2 | P3 | Out | D | std(D) | Z | BABA | ABBA |
| --- | --- | --- | --- | --- | --- | --- | --- | --- |
| BDAP12 | **FLAP79** | CBAP4 | PY21 | 0.905 | 0.054 | 16.72 | 3 | 60 |
| BDAP12 | **FLAP15** | CBAP4 | PY21 | 0.789 | 0.107 | 7.39 | 4 | 34 |
| BDAP12 | **FLAP58** | CBAP4 | PY21 | 0.791 | 0.091 | 8.71 | 7 | 60 |
| BDAP12 | **FLAP64** | CBAP4 | PY21 | 0.803 | 0.080 | 9.97 | 7 | 64 |
| BDAP12 | FLAP23 | **CBAP32** | PY21 | 0.920 | 0.083 | 11.03 | 1 | 24 |
| BDAP12 | FLAP22 | **CBAP11** | PY21 | 0.833 | 0.116 | 7.17 | 2 | 22 |

Table S7. HyDe results for the *Ancylomenes pedersoni* species complex. Only significant evidence of hybridization is reported. Row 1 (highlighted) = results from hybridization detection at the population level. Rows 2-16 = results from hybrization detection at the individual level. Only 1 individual from Bermuda does not appear to be a hybrid between Florida and Caribbean species (BDAP27).

| **P1** | **Hybrid** | **P2** | **Z-score** | **P-value** | **Gamma** |
| --- | --- | --- | --- | --- | --- |
| Florida | Bermuda | Caribbean | 20.62 | ~ 0.00 | 0.48 |
| Florida | BDAP33 | Caribbean | 12.66 | ~ 0.00 | 0.46 |
| Florida | BDAP30 | Caribbean | 22.71 | ~ 0.00 | 0.47 |
| Florida | BDAP22 | Caribbean | 24.65 | ~ 0.00 | 0.48 |
| Florida | BDAP18 | Caribbean | 27.71 | ~ 0.00 | 0.48 |
| Florida | BDAP19 | Caribbean | 23.30 | ~ 0.00 | 0.48 |
| Florida | BDAP27 | Caribbean | -21.56 | 1.0 | 0.51 |
| Florida | BDAP35 | Caribbean | 21.54 | ~ 0.00 | 0.48 |
| Florida | BDAP14 | Caribbean | 20.24 | ~ 0.00 | 0.48 |
| Florida | BDAP15 | Caribbean | 24.32 | ~ 0.00 | 0.48 |
| Florida | BDAP12 | Caribbean | 22.64 | ~ 0.00 | 0.48 |
| Florida | BDAP13 | Caribbean | 26.74 | ~ 0.00 | 0.48 |
| Florida | BDAP29 | Caribbean | 23.01 | ~ 0.00 | 0.47 |
| Florida | BDAP31 | Caribbean | 27.22 | ~ 0.00 | 0.48 |
| Florida | BDAP34 | Caribbean | 21.60 | ~ 0.00 | 0.47 |
| Florida | BDAP28 | Caribbean | 27.00 | ~ 0.00 | 0.48 |

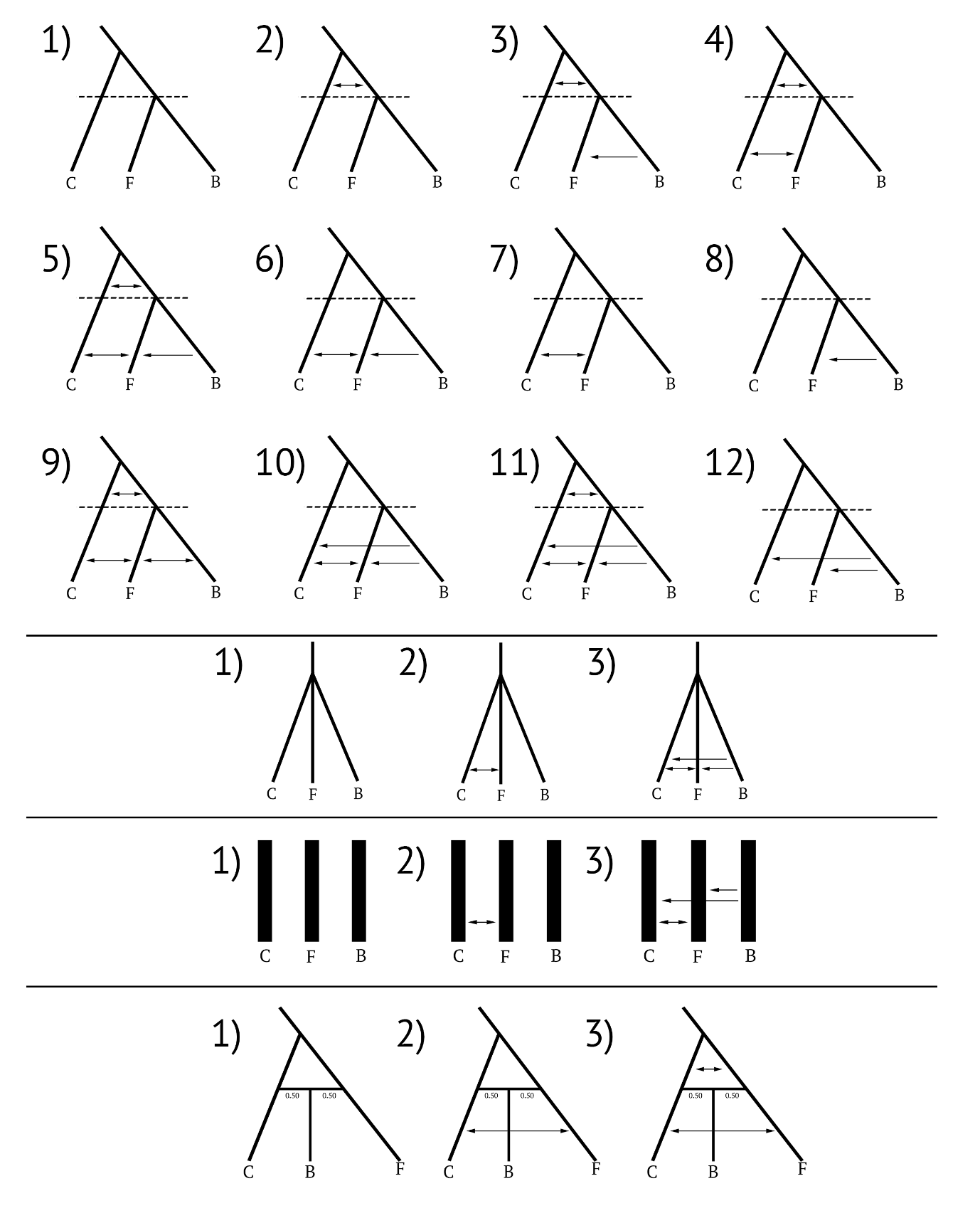

Figure S1. Evolutionary models simulated in *fastsimcoal2* and used for model selection analyses. The first 12 models represent all pairwise migration comparisons made for each species tree topology. Depicted here is the (Caribbean, (Florida, Bermuda)) species tree topology. The same migration comparisons were also made for (Florida, (Caribbean, Bermuda)) and (Bermuda, (Florida, Caribbean)) species tree topologies as well. The next three models depict simultaneous divergence with and without gene flow between each putative species. The third row represents three island models with and without gene flow. The fourth row represents three hybrid speciation models with and without gene flow between Caribbean and Floridian lineages that specify both species provided 50% of the genome to the resulting Bermudan hybrid endemic.
